# A scientific note on “Rapid brood decapping” – a method for assessment of honey bee (*Apis mellifera*) brood infestation with *Tropilaelaps mercedesae*

**DOI:** 10.1101/2024.10.09.616962

**Authors:** Aleksandar Uzunov, Irakli Janashia, Chao Chen, Cecilia Costa, Marin Kovačić

## Abstract

*Tropilaelaps mercedesae* is a parasitic mite species that negatively affects the health of the *Apis mellifera* colonies. Recent reports show that it is spreading westwards, through Central Asia into Europe. Several field and laboratory methods have been proposed to detect *Tropilaelaps* spp. in *A. mellifera* colonies; however, most of them are either laborious, costly or ineffective. Here we introduce a novel method for detecting *T. mercedesae* based on the mite’s characteristic biology (reduced feeding as bee pupae mature, brief dispersal phase on adult bees and agility) and the use of wax strips. Sealed worker brood cells at the development stage of white to purple-eyed pupae are swiftly decapped with wax strips to observe and count surfacing adult mites. Our study shows over 90 % detection efficacy and brood survival, and ease of application. Therefore, we recommend the novel “Rapid brood decapping” method as a reliable tool for detecting and monitoring *T. mercedesae* infestation.

## ARTICLE

*Tropilaelaps mercedesae* (Anderson and Morgan 2007) is one of the 4 currently described species of the *Tropilaelaps* genus that naturally parasitise Asian giant honey bees, *Apis dorsata, A. breviligula and A. laboriosa* (Anderson and Roberts 2013; Chantawannakul et al. 2018). This species also infests the Western honey bee *Apis mellifera* in many Asian countries (Delfinado and Baker 1961; Chantawannakul et al. 2018) and is considered one of its most damaging pests (Rinderer et al. 1994; Dainat et al. 2009; Gao et al. 2021; Han et al. 2024).

Recent reports show that the mite is spreading westwards, through Central Asia into Europe, where it has been found in southwestern Russia and Georgia (Namin et al. 2024; Brandorf et al. 2024; Janashia and Uzunov et al. 2024). Thus, it is evermore crucial for the beekeeping sector to be equipped with reliable methods for rapid detection and effective monitoring.

Several field and laboratory methods, mainly adapted from those used for *Varroa destructor*, have been proposed for the detection of *Tropilaelaps* spp. in *A. mellifera* colonies, such as sampling and analysing adult bees (powder sugar roll, soapy water wash, ethanol wash, CO2) or sealed brood (uncapping with a honey fork or with tweezers/forceps), using the “comb bump”, observing mite fall on bottom boards or analysing hive debris by flotation method in alcohol or by molecular analyses (de Guzman et al. 2017, Kim et al. 2019, Gill et al. 2024). The proposed methods, primarily based on the biology of *V. destructor*, are characterised by being either laborious, costly or ineffective for detecting *Tropilaelaps* spp., and some are even destructive to the colony (Gill 2024; Gill et al. 2024). Gill et al. (2024) found that uncapping infested brood with tweezers, using sticky traps to catch mite drop and rolling adult bees in icing sugar were the most promising methods for detection of *Tropilaelaps* mites.

Here, we introduce a novel method for detecting and monitoring *T. mercedesae* based on the mite’s characteristic biology (reduced feeding as bee pupae mature, brief dispersal phase on adult bees and agility) and the use of wax strips for decapping sealed brood area. Sealed worker brood cells at the development stage of white to purple-eyed pupae (13-17 days from egg laying) are swiftly decapped with wax strips to observe and count surfacing adult mites. To evaluate the method’s detection efficacy, reliability and practicality, we conducted a study in the summer of 2024 in Georgia, using *A. mellifera* colonies (N =30) in apiaries (N = 3) infested with *T. mercedesae*. A comb of sealed worker brood from each colony, with eye colour ranging from white to purple, was decapped using a commercial wax strip (Veet^©^) covering more than 250 cells. The wax strip was evenly applied to the sealed brood area by pressing gently with a soft paper ball for 10 seconds. Once adhered, the strip was immediately and swiftly removed from the comb’s surface. The entire decapped (stripped) brood area was instantly video recorded using a Honor 90 mobile phone camera on an overhead mount for 60 seconds to film and detect emerging mites. Following the video recording, the decapped brood area and the sealed worker brood from the comb’s backside (control cells) were cut out. Care was taken to register any mites leaving the decapped area while cutting brood samples. The comb cuts were wrapped tightly in a white paper towel, inserted into a ziplock sample bag and taken to the laboratory, where they were frozen at – 20°C before inspection of brood infestation.

For each comb sample, *T. mercedesae* infestation was assessed on both sides by examining cell contents using a binocular stereo microscope (Carl Zeiss Stemi 508, magnification x6.3). The control brood cells were carefully uncapped using tweezers (Gill et al. 2024), and the data were used to estimate the correlation with the video observed mites. *V. destructor* infestation was assessed only in the stripped cells. Freezing at -20°C eliminated the risk of counting the same mite twice. Only adult *T. mercedesae* and *V. destructor* mites were considered, identified by the brown pigmentation.

Considering the total number of inspected brood samples (N = 28), an average of 200.9 cells (146 - 241) were checked in the decapped brood and an average of 202.9 previously unopened control cells (151 - 253). Out of 28 samples, 11 (39.3 %) were positive for *T. mercedesae*; of these, 9 samples (81.8 %) were co-infested with *T. mercedesae* and *V. destructor*, and two samples (18.2 %) infested only with *T. mercedesae* (Table 1). The *T. mercedesae* infestation level from the decapped cells, including the video-observed mites (“super control”), averaged 3.05 %. Multiple species infestation of a single cell was detected in only one sample.

**Table 1.**
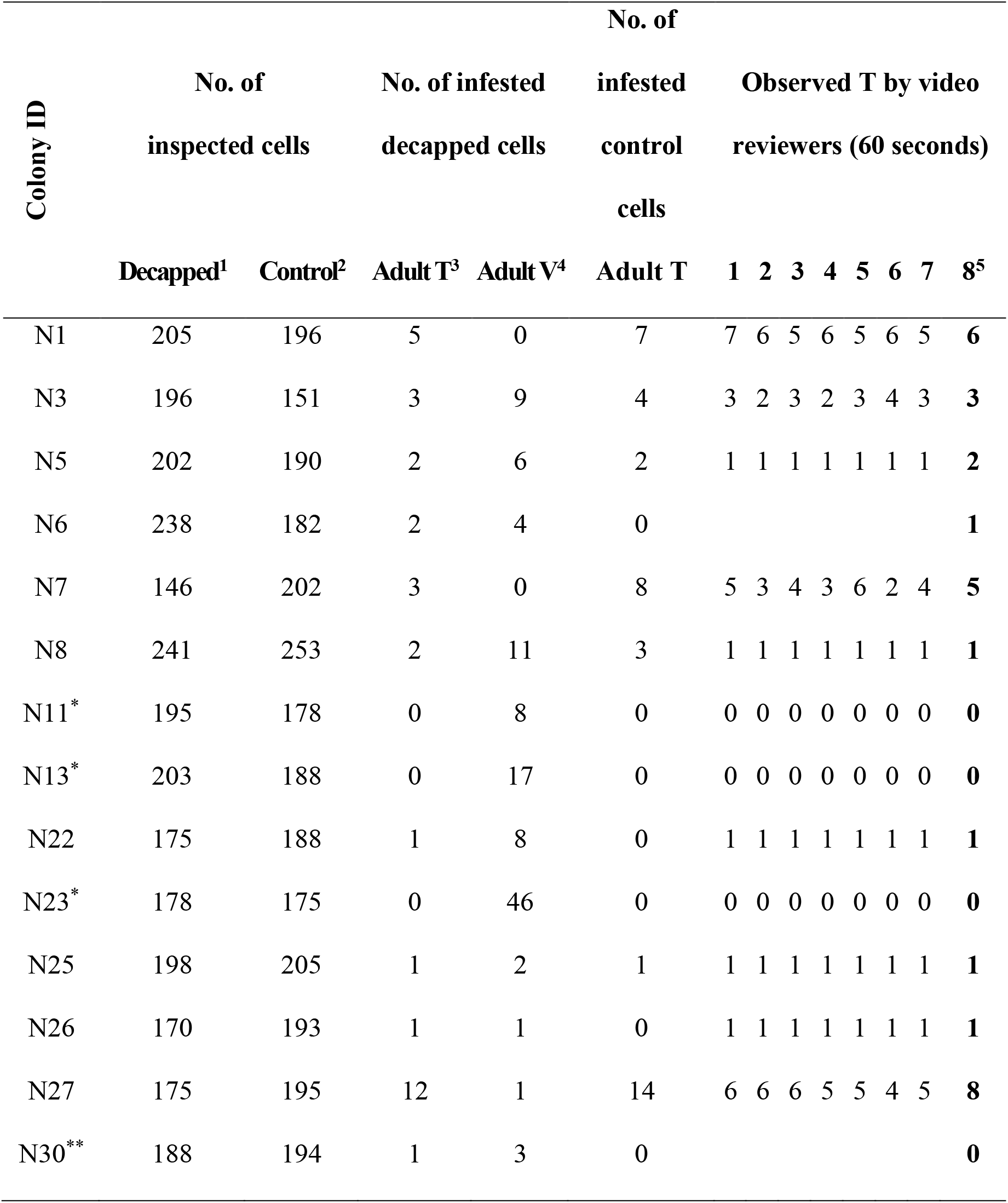
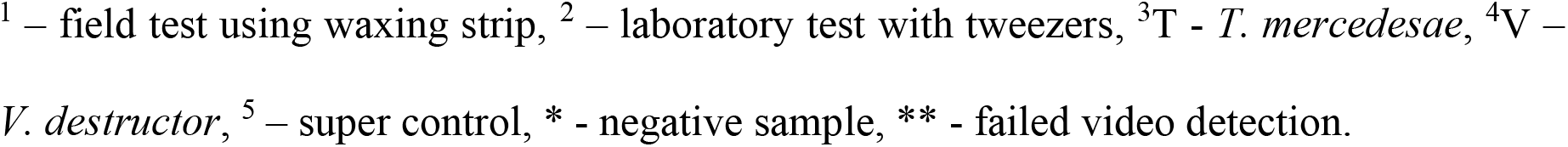
The outcome of the lab inspection of brood and observations of the video recordings post decapping in eleven (N = 11) *T. mercedesae*-positive worker brood samples and three (N = 3) negative samples, which were used as controls in the “blind” video observations.

| Colony ID | No. of |  |  |  |  |  |  |  |  |  |  |  |  |
| --- | --- | --- | --- | --- | --- | --- | --- | --- | --- | --- | --- | --- | --- |
|  | No. of inspected cells |  | No. of infested decapped cells |  | No. of infested control cells |  | Observed T by video reviewers (60 seconds) |  |  |  |  |  |  |
|  | Decapped<sup>1</sup> | Control<sup>2</sup> | Adult T<sup>3</sup> | Adult V<sup>4</sup> | Adult T | 1 | 2 | 3 | 4 | 5 | 6 | 7 | 8<sup>5</sup> |
| N1 | 205 | 196 | 5 | 0 | 7 | 7 | 6 | 5 | 6 | 5 | 6 | 5 | 6 |
| N3 | 196 | 151 | 3 | 9 | 4 | 3 | 2 | 3 | 2 | 3 | 4 | 3 | 3 |
| N5 | 202 | 190 | 2 | 6 | 2 | 1 | 1 | 1 | 1 | 1 | 1 | 1 | 2 |
| N6 | 238 | 182 | 2 | 4 | 0 |  |  |  |  |  |  |  | 1 |
| N7 | 146 | 202 | 3 | 0 | 8 | 5 | 3 | 4 | 3 | 6 | 2 | 4 | 5 |
| N8 | 241 | 253 | 2 | 11 | 3 | 1 | 1 | 1 | 1 | 1 | 1 | 1 | 1 |
| N11* | 195 | 178 | 0 | 8 | 0 | 0 | 0 | 0 | 0 | 0 | 0 | 0 | 0 |
| N13* | 203 | 188 | 0 | 17 | 0 | 0 | 0 | 0 | 0 | 0 | 0 | 0 | 0 |
| N22 | 175 | 188 | 1 | 8 | 0 | 1 | 1 | 1 | 1 | 1 | 1 | 1 | 1 |
| N23* | 178 | 175 | 0 | 46 | 0 | 0 | 0 | 0 | 0 | 0 | 0 | 0 | 0 |
| N25 | 198 | 205 | 1 | 2 | 1 | 1 | 1 | 1 | 1 | 1 | 1 | 1 | 1 |
| N26 | 170 | 193 | 1 | 1 | 0 | 1 | 1 | 1 | 1 | 1 | 1 | 1 | 1 |
| N27 | 175 | 195 | 12 | 1 | 14 | 6 | 6 | 6 | 5 | 5 | 4 | 5 | 8 |
| N30** | 188 | 194 | 1 | 3 | 0 |  |  |  |  |  |  |  | 0 |

<sup>1</sup> – field test using waxing strip, <sup>2</sup> – laboratory test with tweezers, <sup>3</sup>T - *T. mercedesae*, <sup>4</sup>V – *V. destructor*, <sup>5</sup> – super control, \* - negative sample, \*\* - failed video detection.

The video recordings were observed by eight (N = 8) persons: an accurate and repeated observation of each recording was conducted by the most experienced person (N = 1, Table 1 - super control), yielding the ultimate number of observable mites. Then, seven (N = 7) persons “blindly” checked positive (N = 9) and negative (N = 3) samples. The data was analysed assuming that each *T. mercedesae* mite observed emerged from a single cell. While this approach simplifies the analysis, it is important to acknowledge that multiple mites may have originated from the same cell. *T. mercedesae* mites move very fast, and identifying the exact cell of exit is challenging. Thus, for consistency in data interpretation, we proceeded with the one mite per cell assumption.

The video observations detected *T. mercedesae* in ten (N = 10) out of eleven (N = 11), subsequently laboratory-confirmed positive samples. A significant positive correlation (r = 0.89, Figure 1, A) was found between the number of mites detected during 60 seconds post-decapping video recordings and the total number of mites in the decapped cells (excluding video-observed mites) from all positive samples. We also found a significant positive correlation (r = 0.96, Figure 1, B) between the number of video-detected mites and the number of mites from the control cells.

**Figure 1.**
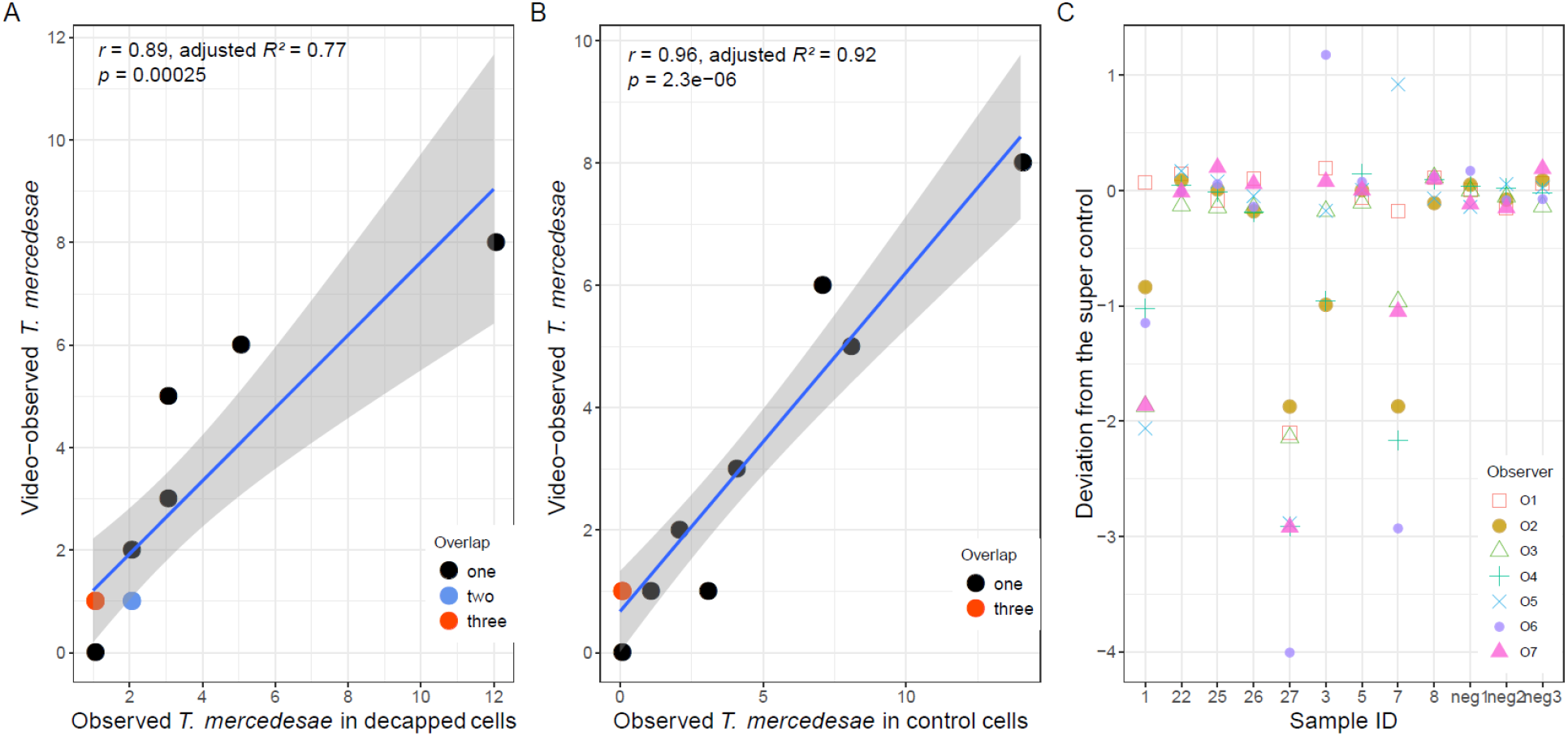
Scatter-plot charts presenting: A. the correlation between the number of video observed (60 sec) mites and the total number of mites for the positive samples (decapped brood cells, excluding the mites from the video observations), B. correlation for the number of video observed mites and the mites from the control side for the positive samples, C. video observers’ deviations from the super control values (Y axis, 0 mark). The grey area indicates 95% confidence interval.

To evaluate the reliability of the observations, the variation between the different observers was analysed (Table 1, excluding samples N6 and N30). All observers successfully distinguished and identified all *T. mercedesae* positive (N = 9) and negative (N = 3) samples. In addition, there was no significant difference (F(7, 88) = 0.111, p > 0.001, One-way ANOVA) in the number of observed mites between observers, indicating that beekeepers and honey bee experts can effectively use this method with different levels of experience (Figure 1, C). No *V. destructor* mites were detected, thus demonstrating the method’s unsuitability in detecting this mite species.

For video observations, simple equipment (monitor and PC) is needed. Due to the high mobility of *T. mercedesae* mites within the first 10 seconds after uncapping, the video review can be improved with particular focus during that time interval. In an average of 19.8 seconds (median 25, SD 12.3, 1 to 55 seconds), the observers reported the definitive number of *T. mercedesae* from the recorded videos. Nevertheless, to ensure accuracy and avoid double counting, we recommend a 30-second observation period. Additionally, assessing the uncapped brood area in two segments/halves may improve detection accuracy in cases of higher infestation.

Additionally, we checked for brood survival 24 hours after the application of the “Rapid brood decapping” method on 29 *T. mercedesae* uninfested *A. mellifera* colonies in Italy (N = 6), Croatia (N = 13) and Georgia (N = 10). The method was found to have low adverse effects on brood survival, with an average of 93.8 % brood survival (SD 10.7, 54.4 to 100 %). Siceanu (1997) reported a similar post-decapping survival (emerging bees) rate of 97.4% using waxed cloth decapping as a *V. destructor* control method. Variations in our study may have been due to differences in strip quality and/or in brood age (e.g. prepupae present in the treated area).

Based on supporting evidence of over 90 % detection efficacy and brood survival, together with ease of application, we recommend the novel “Rapid brood decapping” method as a reliable tool for detecting and monitoring *T. mercedesae* infestation. This method is suitable for beekeeping and research settings, being less invasive and stressful for colonies compared to other existing methods, cost-effective, and quick (around 10 minutes per colony, including colony manipulation and video reviewing). Furthermore, as many pupae are uncovered, this method can also be used to assess covert forms of brood diseases. The step-by-step procedure of the method application is photo and video visualised and described in Uzunov et al. (unpubl. data). We also see a potential for further enhancement of the method through machine learning and mobile device integration. Given the biological similarities among *Tropilaelaps* species, it is likely that the method can be applied across the genus. Further research is needed to validate and refine this method which promises to be a key tool for monitoring and managing *T. mercedesae*’s westward spread.

## ACKNOWLEDGEMENTS

Credit is due to Adrian Siceanu from Romania and Ralph Büchler from Germany for previously suggesting and using this method for Varroa monitoring and control. We thank the students Ruiyi Cheng (程 瑞益), Qi Xu (徐 齐) and Yufei Zou (邹禹妃) and the Georgian beekeepers for their support.

## AUTHOR CONTRIBUTION

AU conceptualised and designed the study. IJ did laboratory analyses. IJ, CeC, and MK did the fieldwork. AU, IJ, ChC, CeC, MK and the students did video reviews. All authors contributed to data processing and interpretation. AU wrote the first draft of the manuscript, and all authors commented on previous versions. All authors read and approved the final manuscript.

## DATA AVAILABILITY

The datasets generated during and/or analysed during the current study are available from the corresponding author upon reasonable request.

## CODE AVAILABILITY

Not applicable

## DECLARATIONS

### Ethics approval

No approval of research ethics committees was required to accomplish the goals of this study because experimental work was conducted with an unregulated invertebrate species.

### Consent to participate

Not applicable.

### Consent for publication

Not applicable.

